# Myocardial *Notch-Rbpj* deletion does not affect heart development or function

**DOI:** 10.1101/393348

**Authors:** Alejandro Salguero-Jiménez, Joaquim Grego-Bessa, Gaetano D’Amato, Luis J. Jiménez-Borreguero, José Luis de la Pompa

## Abstract

During vertebrate cardiac development NOTCH signaling activity in the endocardium is essential for the crosstalk between endocardium and myocardium that initiates ventricular trabeculation and valve primordium formation. This crosstalk leads later to the maturation and compaction of the ventricular chambers and the morphogenesis of the cardiac valves, and its alteration may lead to disease. Although endocardial NOTCH signaling has been shown to be crucial for heart development, its physiological role in the myocardium has not been clearly established. Here we have used a genetic strategy to evaluate the role of NOTCH in myocardial development. We have inactivated the unique and ubiquitous NOTCH effector RBPJ in the early cardiomyocytes progenitors, and examined its consequences in cardiac development and function. Our results demonstrate that mice with *cTnT-Cre*-mediated myocardial-specific deletion of *Rbpj* develop to term, with homozygous mutant animals showing normal expression of cardiac development markers, and normal adult heart function. Similar observations have been obtained after *Notch1* deletion with *cTnT-Cre*. We have also deleted *Rbpj* in both myocardial and endocardial progenitor cells, using the *Nkx2.5-Cre* driver, resulting in ventricular septal defect (VSD), double outlet right ventricle (DORV), and bicuspid aortic valve (BAV), due to NOTCH signaling abrogation in the endocardium of cardiac valves territories. Our data demonstrate that NOTCH-RBPJ inactivation in the myocardium does not affect heart development or adult cardiac function.

## Introduction

The heart is the first organ to form and function during vertebrate development. At embryonic day 7.0 (E7.0) in the mouse, cardiac progenitor cells, migrating from the primitive streak, reach the head folds on either side of the midline (1) and by E8.0, fuse and form the primitive heart tube (2). The heart tube consists internally of the endocardium, that is separated from the primitive myocardium by an extracellular matrix termed cardiac jelly (3). The NOTCH signaling pathway is crucial for the endocardial-myocardial interactions that regulate the patterning, growth and differentiation of chamber and non-chamber tissues that will develop from E8.5 onwards (4–8). The main components of the pathway are the single-pass transmembrane NOTCH receptors (NOTCH1-4 in mammals) that interact with membrane-bound ligands of the JAGGED (JAG1 and JAG2) and DELTA families (DELTA LIKE1, 3 and 4), expressed in neighboring cells (9, 10). Ligand-receptor interactions leads to three consecutive cleavage events that generate the NOTCH intracellular domain (NICD), which can translocate to the nucleus of the signaling-receiving cell (11). In the nucleus, NICD binds directly to the DNA-binding protein CSL (CBF1/RBPJ/Su(H)/Lag1) (12) and recruits the co-activator Mastermind-like (13, 14). In the absence of N1ICD, ubiquitously expressed RBPJ (recombination signal binding protein for immunoglobulin kappa J region) may act as a transcriptional repressor (15). The best characterized NOTCH targets in the heart are the HEY family of basic helix–loop–helix (bHLH) transcriptional repressors (16), although various other cardiac-specific targets have been described (4, 17–19). Functional studies in *Xenopus* or in *Rbpj*-targeted mouse embryonic stem cells have shown that NOTCH suppresses cardiomiogenesis (20, 21), although studies with targeted mutant mice have demonstrated an essential requirement for NOTCH in cardiac development only after heart tube formation (around E8.5) (22, 23).

One of the first sign of cardiac chamber development is the appearance of trabeculae at E9.0-9.5 (24). Trabeculae are myocardial protrusions covered by endocardium that grow towards the ventricular lumen, and serve to facilitate oxygen exchange and nourishment between the blood and the developing heart. The ligand DLL4 and the active NOTCH1 receptor are expressed in the endocardium prior to the onset of trabeculation (4, 25). DLL4-NOTCH1 signaling is reflected by the endocardial expression of the CBF:H2B-Venus transgenic NOTCH reporter in mice (17). Conditional inactivation of *Dll4, Notch1* or *Rbpj* in the endocardium, results in very similar phenotypes (more severe in *Rbpj* mutants) consisting of ventricular hypoplasia and impaired trabeculation (4, 17, 26), while myocardial deletion of *Jag1* does not affect trabeculation (17). Later, NOTCH1 signaling in the endocardium is activated in a temporal sequence from the myocardium by the JAGGED1/2 ligands in a MIB1-dependent manner, to sustain ventricular compaction and maturation (17, 27). Conditional myocardial deletion of *Mib1* or combined inactivation of *Jag1* and *Jag2* abrogates endocardial NOTCH activity, and leads to abnormally thin compact myocardium and large and non-compacted trabeculae, a phenotype strongly reminiscent of a cardiomyopathy termed left ventricular non-compaction (LVNC) (17, 27). Thus, endocardial NOTCH signaling and its downstream effectors are essential for the endocardium-to-myocardium signaling that regulates chamber patterning and growth (28–30). Endocardial NOTCH activity is also crucial for development and morphogenesis of the cardiac valves (5, 18, 29, 31–37). NOTCH receptors or transgenic reporter lines are expressed in the endocardium, coronary vessels endothelium and smooth muscle cells (17, 25, 38, 39). There is not clear evidence of endogenous NOTCH expression or activity in the embryonic myocardium, and despite elegant ectopic expression experiments have reported a function for NOTCH in cardiomyocytes (40, 41), a physiological role for NOTCH in the developing myocardium has not been clearly demonstrated *in vivo*.

To address this question, we have conditionally inactivated the NOTCH effector RBPJ in the myocardium using two early-acting myocardial drivers, and examined its consequences in heart development. We find that *cTnT-Cre* mediated myocardial-specific deletion of *Rbpj* does not affect NOTCH signaling in the endocardium, heart development or adult heart function. In contrast, while *Nkx2.5-Cre* mediated *Rbpj* inactivation in the myocardium does not affect cardiac development and structure, *Rbpj* inactivation in valve endocardial cells disrupts valve morphogenesis. Our data demonstrate that myocardial NOTCH-RBPJ is not required for cardiac development or function, reinforcing the notion that physiological NOTCH-RBPJ signaling occurs in the endocardium, endothelium and smooth muscle cells of the developing heart.

## Results and Discussion

We first compared expression of the NOTCH effector RBPJ to the pattern of NOTCH activation in the E12.5 heart. RBPJ was widely expressed in the nucleus of endocardial, myocardial and epicardial cells (Fig 1a-a’), while NOTCH1 activity was restricted to the endocardium (Fig 1b-b’). Expression of the NOTCH transgenic reporter *CBF:H2B-Venus* in endocardial cells indicated that NOTCH activity was restricted to the endocardium (Fig 1c-c’).

**Fig 1.**
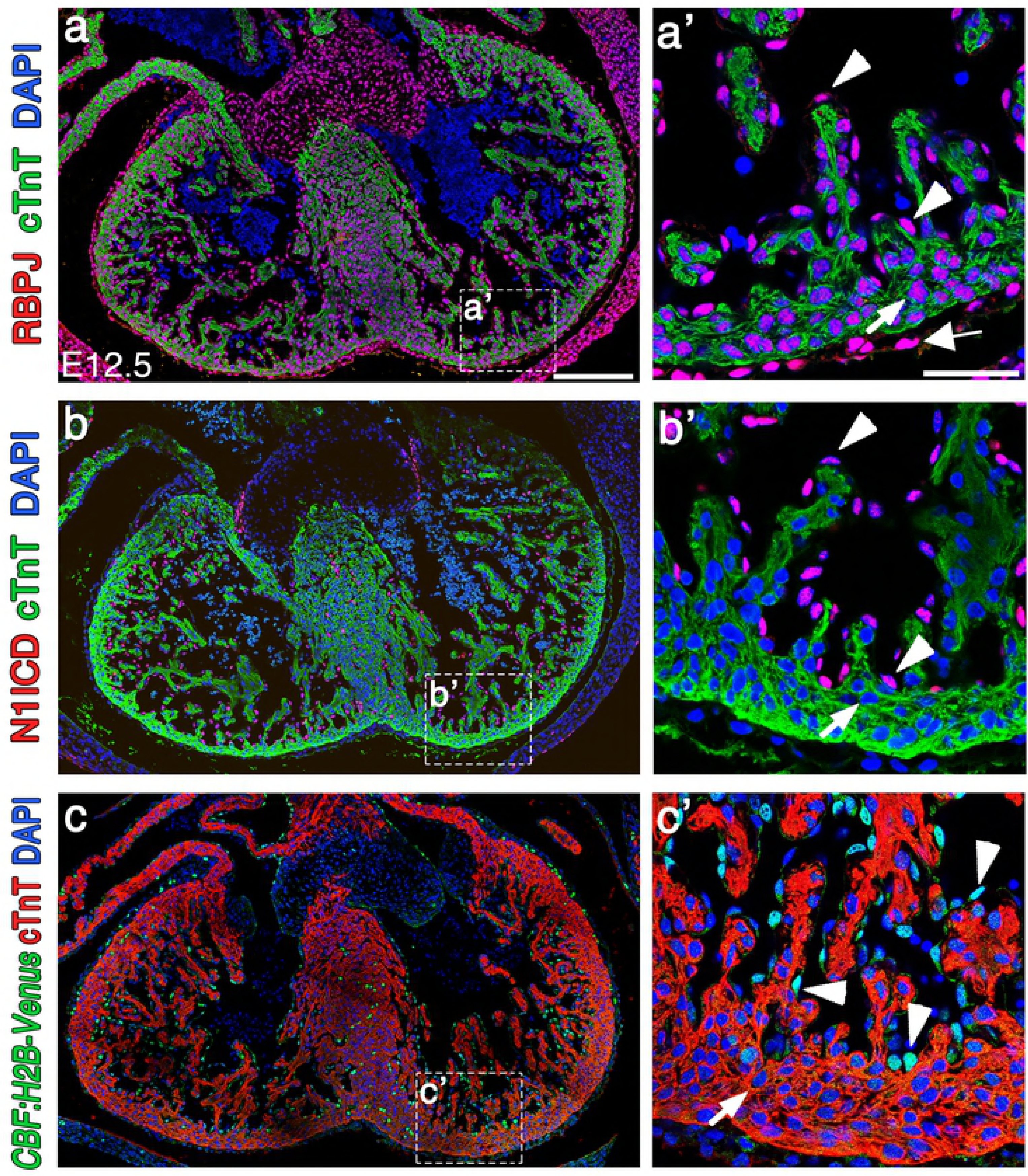
RBPJ is ubiquitously expressed in the nucleus of cardiac cells, while NOTCH activity is restricted to the endocardium during cardiac development. (a-a’) RBPJ (red) and (b-b’) N1ICD (red) nuclear immunostaining in wild type (WT) E12.5 cardiac sections. (c-c’) *CBF:H2B-Venus* reporter line expression (green) in E12.5 cardiac sections. The myocardium is cTnT-counterstained (green in a-b’, red in c-c’). White arrows indicate cardiomyocytes, white arrowheads point to endocardial cells, and the thick arrow in (a’) indicates epicardial RBPJ expression. Note that cardiomyocytes do not express CBF:H2B-Venus (c,c’). Scale bars: a-c, 200μm, a’-c’, 50μm.

We then generated myocardial-specific conditional mutants by breeding *Rbpj^flox/flox^* mice (42) with *cTnT-Cre^tg/+^* mice, which express the CRE recombinase specifically in cardiomyocytes from E8.0 onwards (43). At E16.5, the heart of *Rbpj^flox^;cTnT-Cre* (*Rbpj^flox/flox^;cTnT^Cre/+^*) embryos was indistinguishable from control (*Rbpj^flox/flox^;cTnT^+/+^*) littermates (Fig 2a-b’), and compact and trabecular myocardium thickness was similar in both genotypes (Fig 2c). Immunostaining confirmed full myocardial RBPJ deletion in E16.5 *Rbpj^flox^;cTnT-Cre* embryos (Fig 2d-e’’). Thus, while in control embryos RBPJ was found in the nucleus of both endocardium and myocardium (Fig 2d-d’’), it was not detected in cardiomyocytes of *Rbpj^flox^;cTnT-Cre* embryos while endocardial RBPJ expression was normal (Fig 2e-e’’).

**Fig 2.**
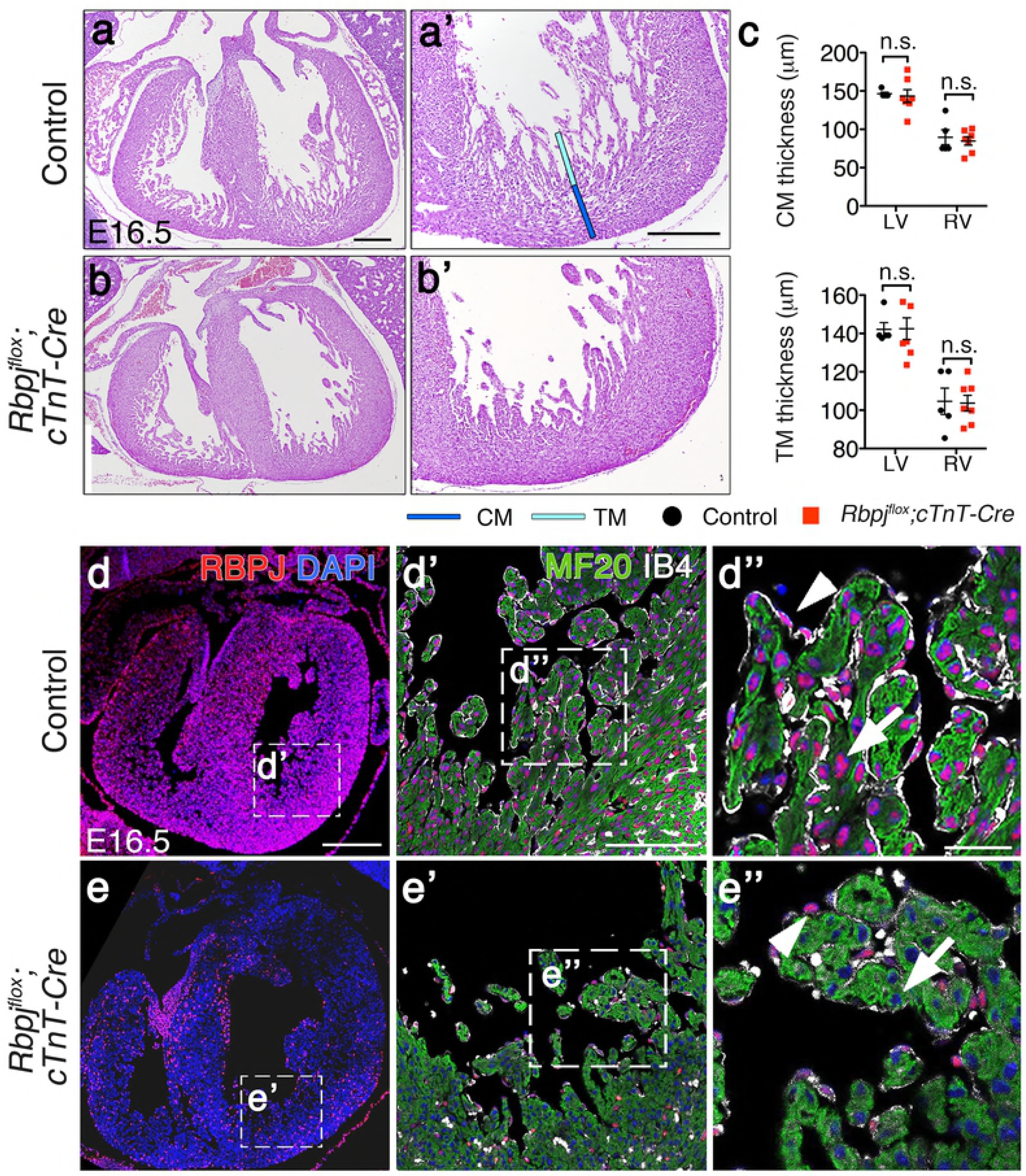
Myocardial *Rbpj* deletion does not affect ventricular development and structure. (a-b’) Hematoxylin and eosin (H&E) staining of heart sections from control and *Rbpj^flox^;cTnT-Cre* E16.5 embryos. (c) Quantification of compact myocardium (CM) and trabecular myocardium (TM) thickness in E16.5 control and *Rbpj^flox^;cTnT-Cre* embryos. LV CM control = 146.4 ± 2.1 μm, LV CM mutant = 143.4 ± 8.4 μm, RV CM control = 89.6 ± 9.8 μm, RV CM mutant = 84.7 ± 5.5 μm, LV TM control = 142.2 ± 3.5 μm, LV TM mutant = 142.5 ± 5.7 μm, RV TM control = 117.0 ± 5.5 μm, RV TM mutant = 115.5 ± 7.6 μm (Data are mean ± s.e.m; *n* = 5 control embryos and *n* = 7 mutant embryos; *P*<0.05 by Student’s *t*-test; n.s., not significant). (d-e’’) RBPJ (red) immunostaining of control and *Rbpj^flox^;cTnT-Cre* E16.5 cardiac sections, myosin heavy chain (MF20, green), and isolectin B4 (IB4, white). White arrows indicate cardiomyocytes; white arrowheads point to endocardial cells. Scale bars: 200μm in a,a’,d; 100μm in d’; 25μm in d’’.

Genetic manipulation of NOTCH elements leading to signal inactivation in the endocardium, disrupts myocardial patterning and chamber maturation (17, 27). We analyzed if ventricular patterning was affected after deletion of RBPJ in the myocardium. E16.5 *Rbpj^flox^;cTnT-Cre* embryos showed normal expression of both compact (*Hey2*) (44–46) and trabecular myocardial markers (*Bmp10* (47), *Cx40* (48)) (Fig 3a-f), indicating that myocardial patterning did not require myocardial RBPJ.

**Fig 3.**
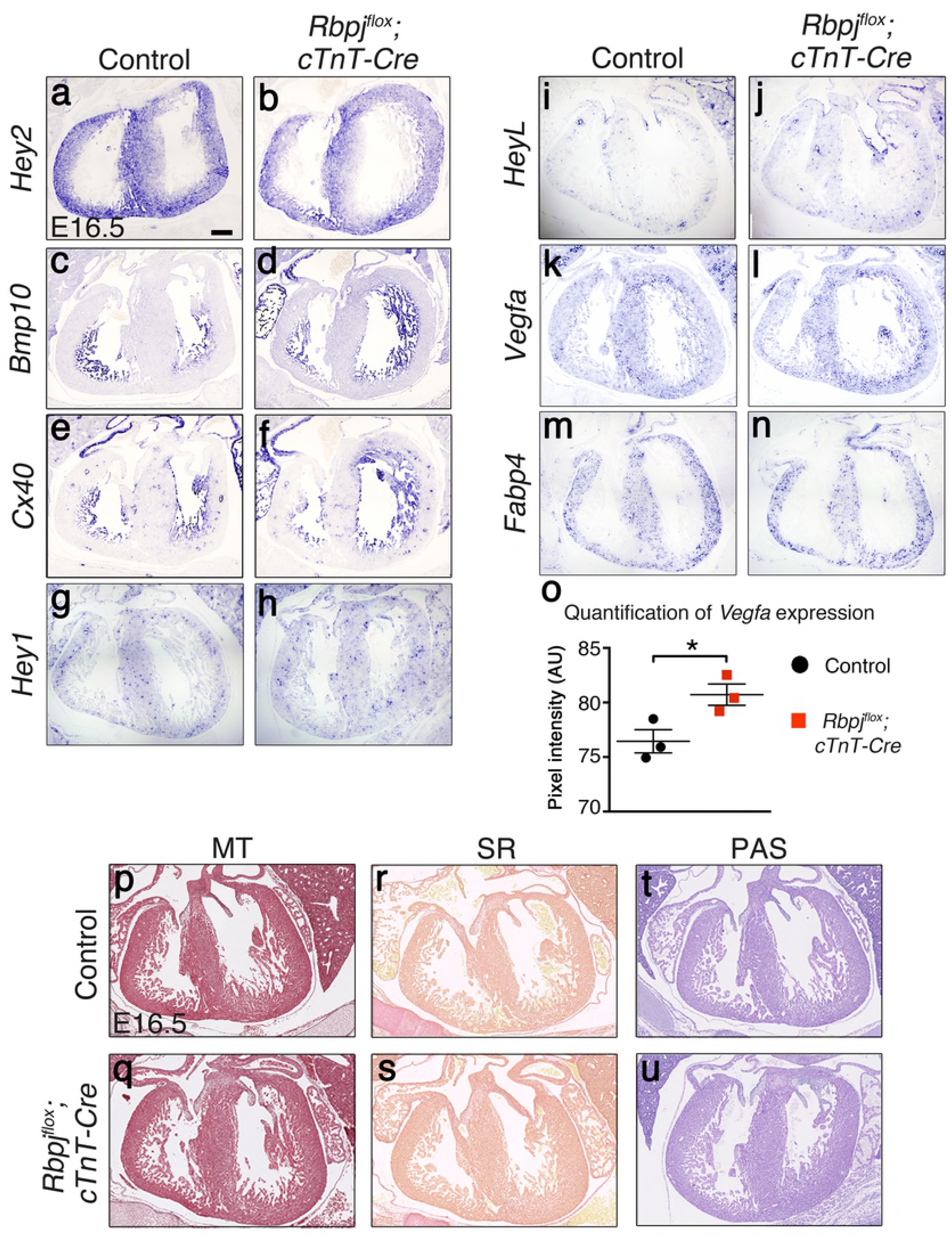
Expression pattern of compact and trabecular myocardium markers, NOTCH target genes, and fibrosis marker staining is normal in *Rbpj^flox^;cTnT-Cre* embryos, while *Vegf* is increased. *In situ* hybridization (ISH) of *Hey2* (a-b), *Bmp10* (c-d)*, Cx40* (e-f)*, Hey1* (g-h), *HeyL* (i-j), *Vegfa* (k-l) and *Fabp4* (m-n) in E16.5 *Rbpj^flox^;cTnT-Cre* and control hearts. (o) *Vegfa* expression quantification from *Vegfa* ISH. E16.5 *Rbpj^flox^;cTnT-Cre* and control cardiac sections stained with Masson’s Trichrome (MT), (p-q); Periodic acid-Schiff (PAS), (r-s), and Sirius Red (SR), (t-u). Scale bar is 200μm.

Canonical NOTCH signaling requires NICD binding to RBPJ in the nucleus to activate target genes expression (11). The Notch target genes *Hey1* and *HeyL* are expressed in both endocardium and coronaries endothelium of E16.5 wild type embryos (Fig 3g-j). The endothelial-endocardial pattern of *Hey1* and *HeyL* expression was maintained E16.5 *Rbpj^flox^;cTnT-Cre* embryos, indicating that *Rbpj* deletion in the myocardium did not affect Notch targets expression in the heart. These results are consistent with the data showing that NOTCH activity in the heart is restricted to endocardium (Figure 1c,c’) and coronaries endothelium, and that physiological NOTCH activity does not occur in the embryonic myocardium (4, 17, 25).

A previous report in which *Rbpj* was inactivated in the myocardium using the *αMhc-Cre* driver (49) showed that myocardial RBPJ represses hypoxia-inducible factors (HIFs) to negatively regulate *Vegfa* expression in a NOTCH-independent manner (50). *In situ* hybridization of *Vegfa* in E16.5 *Rbpj^flox^;cTnT-Cre* embryos showed a small but significant increase of *Vegfa* transcription in the ventricular wall of mutant embryos (Fig 3k-l, o), supporting previous observations (50). VEGFA positively regulates the formation of blood vessels in the ventricles (51). Thus, we analyzed the expression of the coronary vessels marker *Fabp4* (52) and observed a similar pattern and intensity in E16.5 *Rbpj^flox^;cTnT-Cre* and control embryos (Fig 3m-n) suggesting that coronaries development was normal. Thus, in agreement with Díaz-Trelles et al., these results revealed a NOTCH-independent role for RBPJ in the negative regulation of *Vegfa* in the myocardium.

Cardiomyopathies may result in the appearance of fibrosis and accumulation of collagen fibers in the myocardium, due to defective vascularization or loss of metabolic homeostasis (53). We performed Masson’s Trichrome and Sirius Red staining in E16.5 *Rbpj^flox^;cTnT-Cre* mutant hearts, and found no signs of fibrosis (Fig. 3p-s). Periodic Acid-Schiff (PAS) staining detects glycogen accumulation that could be induced by inflammation, but PAS staining was relatively normal in E16.5 *Rbpj^flox^;cTnT-Cre* hearts (Fig 3t-u). These results indicated that *Rbpj* deletion in the embryonic myocardium does not affect myocardial fetal development.

*Rbpj^flox^;cTnT-Cre* mutant mice reached adulthood in similar proportions than control littermates. Genotyping of neonatal and adult litters showed that all genotypes appeared at the expected Mendelian proportions (Table 1), indicating that RBPJ loss in the myocardium did not compromise postnatal viability. Morphological analysis of 6-month old *Rbpj^flox^;cTnT-Cre* adults revealed normal heart structure compared to control animals (Fig 4a,b). In order to detect potential physiological impairments in the heart, we analyzed cardiac function by echocardiography (Fig 4e). Ejection fraction (EF%) and fractional shortening (FS%) were similar in wild type and mutant mice (Fig 4e). The diastolic function, indicated by the E/A ratio (ratio of early diastolic velocity to atrial velocity) (54), was also normal. Physiological measurements indicate that ventricular volumetric and mass parameters were normal compared to control mice. Overall, the echocardiography study indicates that myocardial *Rbpj* inactivation does not affect postnatal heart growth and adult myocardial function. We confirmed these results by inactivating *Notch1* with the *cTnT-Cre* driver. Six-month old *Notch1^flox^;cTnT-Cre* adult hearts did not show any morphological phenotypes compared to control animal (Fig. 4c,d). In terms of heart function, *Notch1^flox^;cTnT-Cre* mice exhibited a slightly better cardiac performance with a minor but significantly increased EF% and FS% compared to control mice (Fig 4f). The diastolic function was normal, as it was in *Rbpj^flox^;cTnT-Cre*.

**Table 1.**
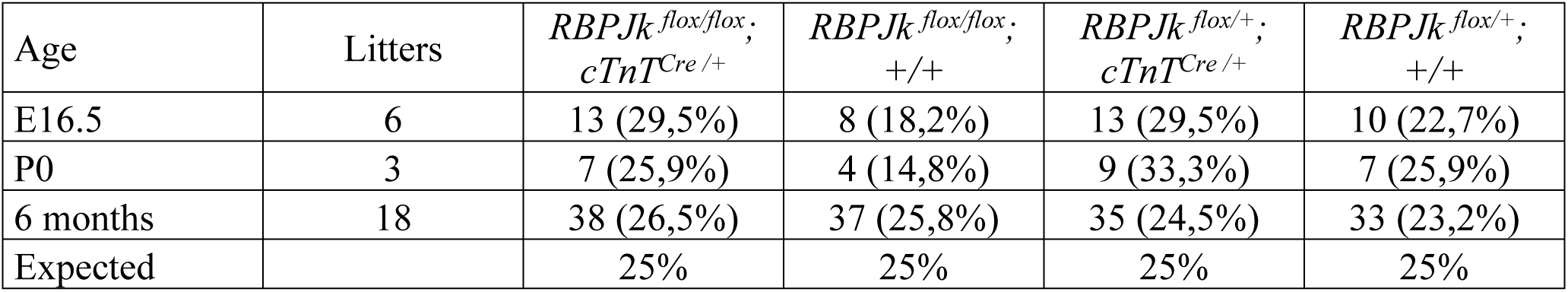
Myocardium-specific *Rbpj^flox^* mutants are viable and reach adulthood. Distribution of the different genetic combinations resulted from the intercross of *Rbpj^flox/+^*; *cTnT^Cre/+^*; males with *Rbpj^flox/flox^* females compared to the expected Mendelian proportions.

**Fig 4.**
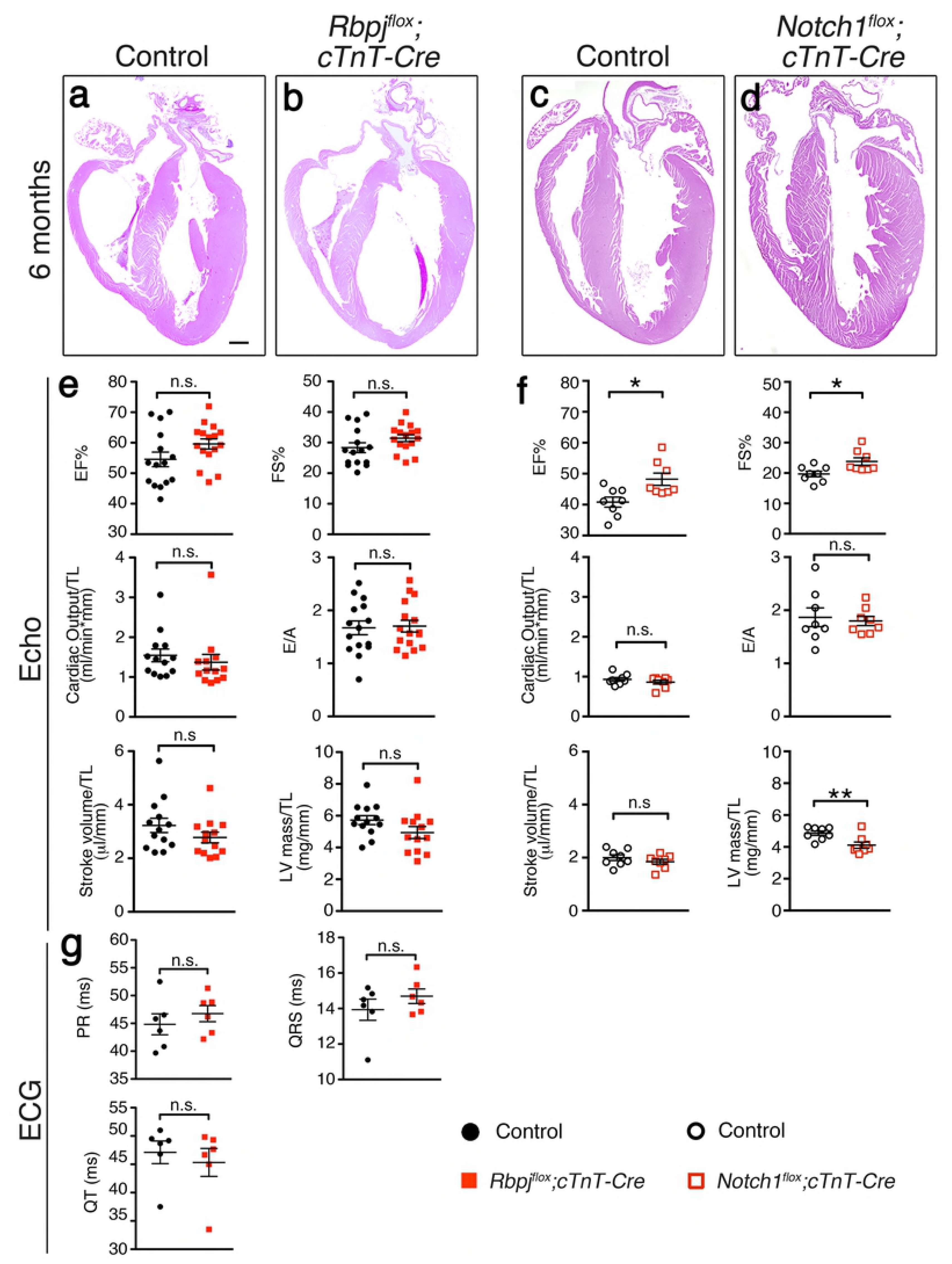
Cardiac structure and function are preserved in both *Rbpj^flox^;cTnT-Cre* and *Notch1^flox^;cTnT-Cre* mice. a-d) H&E staining of cardiac sections from six months-old *Rbpj^flox^ cTnT-Cre* and *Notch1^flox^;cTnT-Cre* mice and their control littermates. (e,f) Echocardiography analysis of six months-old *Rbpj^flox^;cTnT-Cre* and *Notch1^flox^;cTnT-Cre* mice. For *Rbpj^flox^; cTnT-Cre:* EF% control = 54.5 ± 2.4, EF% mutant= 59.6 ± 1.7, FS% control = 28.3 ± 1.6, FS% mutant = 31.4 ± 1.1, E/A control = 1.7 ± 0.1, E/A mutant = 1.7 ±0.1, Cardiac Output/TL control = 1.5 ± 0.2, Cardiac Output/TL mutant = 1.4 ± 0.2, Stroke volume/TL control = 3.2 ± 0.3, Stroke volume/TL mutant = 2.8 ± 0.2, LV mass/TL control = 5.7 ± 0.3, LV mass/TL mutant = 4.9 ± 0.4. *Notch1^flox^;cTnT-Cre:* EF% control = 40.8 ± 1.6, EF% mutant = 48.21 ± 2.0, FS% control = 19.7 ± 0.9, FS% mutant = 23.8 ± 1.2, E/A control = 1.9 ± 0.2, E/A mutant = 1.8 ± 0.4, Cardiac Output/TL control = 0.9 ± 0.1, Cardiac Output/TL mutant = 0.9 ± 0.1, Stroke Volume/TL control = 2.0 ± 0.1, Stroke Volume/TL mutant = 1.8 ± 0.1, LV mass/TL control = 4.9 ± 0.1, LV mass/TL mutant = 4.1 ± 0.2. (g) Electrocardiogram analysis of control and *Rbpj^flox^; cTnT-Cre* 6 months old mice. PR control = 44.8 ± 1.9, PR mutant = 46.7 ± 1.4, QRS control = 13.9 ± 0.6, QRS mutant = 14.7 ± 0.4, QT control = 47.1 ± 2.0, QT mutant = 43.3 ± 2.5. Data are mean ± s.e.m. For echo, *n* = 15 *Rbpj^flox^* control mice, *n* = 16 *Rbpj^flox^;cTnT-Cre* mutant mice, *n* = 8 *Notch1^flox^* control mice, *n* = 8 *Notch1^flox^;cTnT-Cre* mutant mice. For ECG, *n* = 6 control mice and *n* = 6 mutant mice. *P*<0.05 by Student’s *t*-test; n.s., not significant; \**P<0.05;* \*\**P*<0.01. EF, ejection fractions; FS, fractional shortening; TL, tibial length; LV, left ventricle. Scale bars is 600μm.

Previous reports suggested that ectopic myocardial Notch signaling directs the differentiation of cardiomyocytes towards specialized conduction cells *in vitro* (41). Although the expression of the ventricular conduction system marker *Cx40* was normal in E16.5 *Rbpj^flox^;cTnT-Cre* mutant embryos (Fig 3e-f), we further analyzed cardiac conduction system activity of *Rbpj^flox^;cTnT-Cre* adult mice. Electrocardiogram showed no significant differences neither in the main intervals PR and QT nor in the QRS complex duration compared to control mice, suggesting that the conduction system is fully functional in *Rbpj^flox^;cTnT-Cre* adult mice (Fig 4g).

To further confirm that myocardial RBPJ is dispensable for cardiac development and function, we used a second early myocardial driver *Nkx2.5-Cre*, active in the myocardium and in a subset of endocardial cells from E7.5 onwards (55). Morphological analysis of E16.5 *Rbpj^flox/flox^;Nkx2.5^Cre/+^* (*Rbpj^flox^;Nkx2.5-Cre*) embryos revealed the presence of a membranous ventricular septal defect (VSD, Fig 5a-b) and dysmorphic valves (Fig 5c-d). Among thirteen *Rbpj^flox^;Nkx2.5-Cre* mutants examined, twelve (92%) showed membranous VSD, twelve (92%) showed double outlet right ventricle (DORV), in which the aorta is connected to the right ventricle instead of to the left one (Fig 5e-f). Seven mutants (54%) had bicuspid aortic valve (BAV), characterized by either right to non-coronary (75% of cases) or right to left (25% of cases) morphology and resulting in a two-leaflet valve instead of the normal three-leaflet valve (Fig 5c-d). *Rbpj^flox^;Nkx2.5-Cre* mutant embryos developed a normal compact and trabecular myocardium layers, with a thickness similar to controls (Fig 5g). *Rbpj^flox^;Nkx2.5-Cre* mutants showed perinatal lethality and died around postnatal day 0 (P0; Table 2).

**Table 2.**
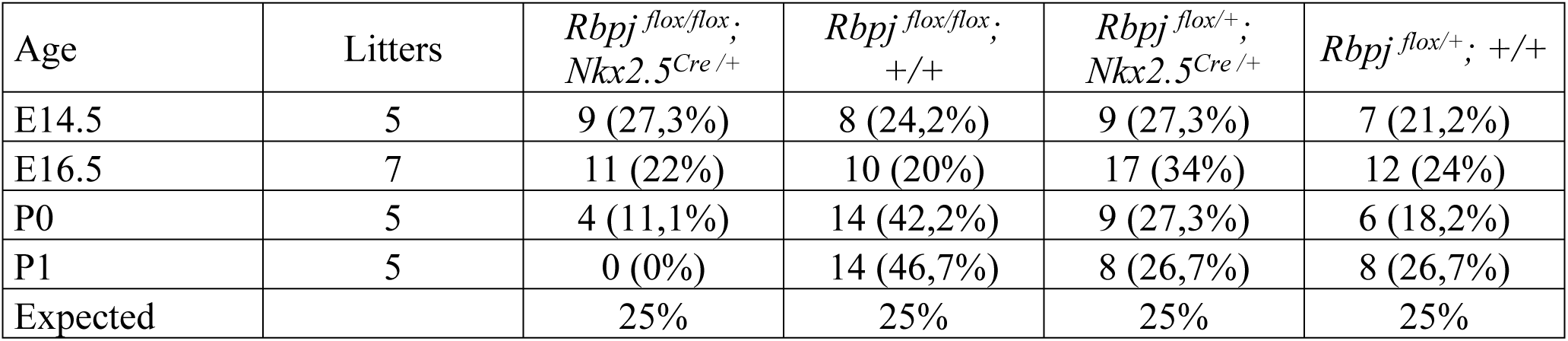
*Rbpj ^flox^; Nkx2.5-Cre* embryos show perinatal lethality. Distribution of the different genetic combinations resulted from the intercross of *Rbpj ^flox/+^; Nkx2.5^Cre^* males with *Rbpj ^flox/flox^* females compared to the expected Mendelian proportions.

| Age | Litters | *Rbpj<sup>flox/flox</sup>; Nkx2.5<sup>Cre</sup> /+* | *Rbpj<sup>flox/flox</sup>; +/+* | *Rbpj<sup>flox/+</sup>; Nkx2.5<sup>Cre</sup> /+* | *Rbpj<sup>flox/+</sup>; +/+* |
| --- | --- | --- | --- | --- | --- |
| E14.5 | 5 | 9 (27,3%) | 8 (24,2%) | 9 (27,3%) | 7 (21,2%) |
| E16.5 | 7 | 11 (22%) | 10 (20%) | 17 (34%) | 12 (24%) |
| P0 | 5 | 4 (11,1%) | 14 (42,2%) | 9 (27,3%) | 6 (18,2%) |
| P1 | 5 | 0 (0%) | 14 (46,7%) | 8 (26,7%) | 8 (26,7%) |
| Expected |  | 25% | 25% | 25% | 25% |

**Figure 5.**
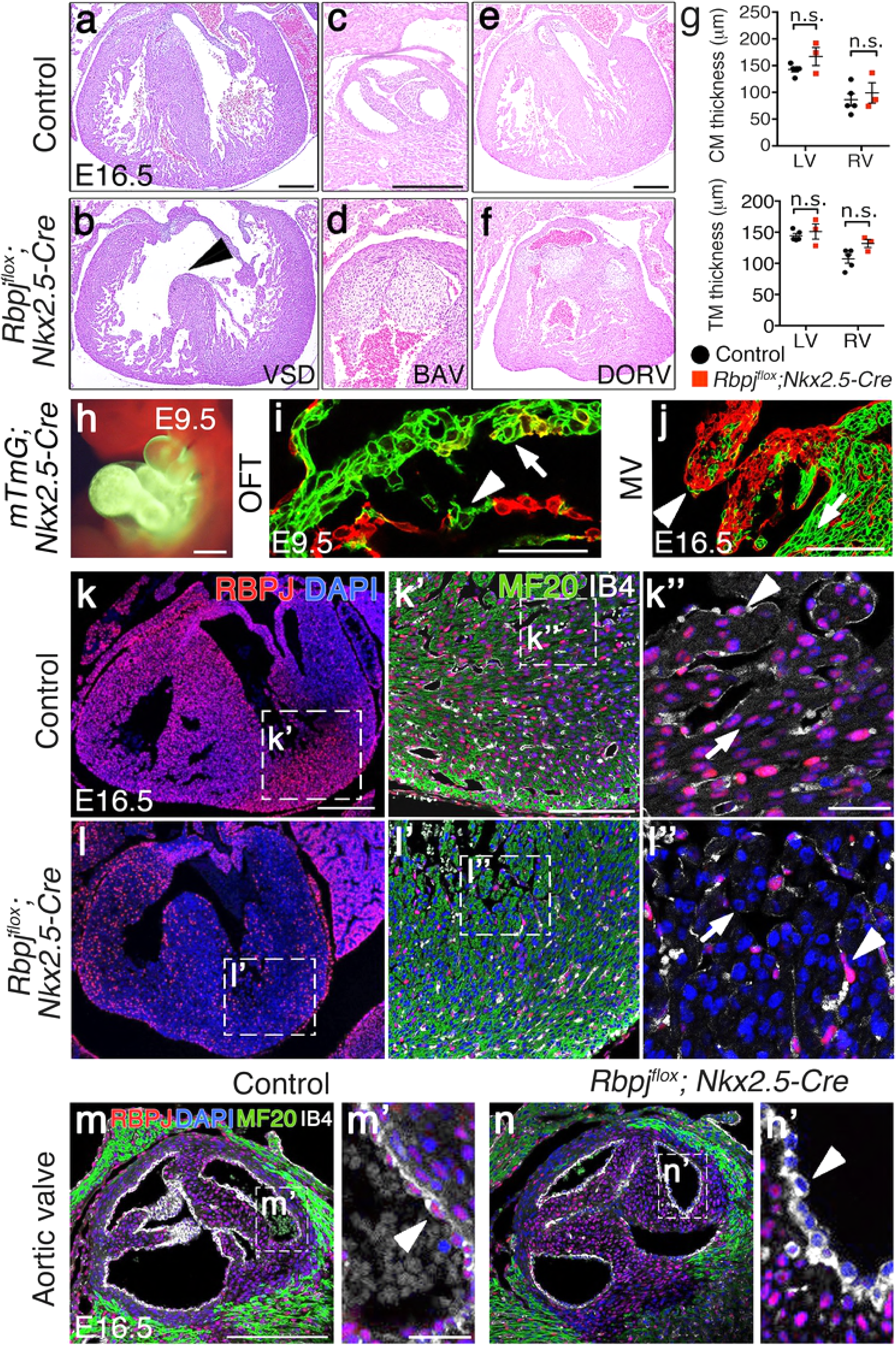
*Nkx2.5-Cre*-mediated *Rbpj* deletion results in ventricular septal defect, bicuspid aortic valve and double outlet right ventricle. (a-f) H&E staining of cardiac sections from E16.5 control and *Rbpj^flox^;Nkx2.5-Cre* embryos showing ventricular septal defect (a,b), bicuspid aortic valve (c,d), and double outlet right ventricle (e,f). (g) Quantification of compact myocardium (CM) and trabecular myocardium (TM) thickness in E16.5 control and *Rbpj^flox^; Nkx2.5-Cre* embryos. LV CM control = 142.8 ± 4.5 μm; LV CM mutant = 167.0 ± 16.8 μm; RV CM control = 86.2 ± 11.4 μm, RV CM mutant = 99.0 ±19.0 μm; LV TM control = 144.3 ± 3.7 μm, LV TM mutant = 151.1 ± 12.2 μm; RV TM control = 116.2 ± 55.5 μm, RV TM mutant = 131.9 ± 6.5 μm (Data are mean ± s.e.m*, n*= 5 control embryos and *n*=3 mutant embryos; *P*<0,05 by Student’s *t*-test; n.s, not significant). (h-j) *Nkx2.5-Cre* lineage tracing analysis using *mTmG* mice show recombination in the entire heart at E9.5 (h), including the outflow track (OFT) endocardium (i). At E16.5, *mTmG;Nkx2.5-Cre* shows partial recombination both at the mitral valve endocardium and in endocardium-derived mesenchyme (j). (k-n’) RBPJ (red), myosin heavy chain (MF20, green) and isolectin B4 (IB4, white) immunostaining in E16.5 control and *Rbpj^flox^;Nkx2.5-Cre* embryos. White arrows indicate cardiomyocytes; white arrowheads point to endocardial cells. Scale bar: 200μm in a-f, k,l; 100μm in h,j,k’,m,n; 50μm in i; 25μm in m’.

To determine the precise contribution of *Nkx2.5*-expressing cells to the developing heart, we took advantage of the *mTmG* system in which following CRE-mediated excision, the *mTomato* transgene is removed so that the *CAG* promoter drives the expression of membrane localized EGFP (56). Lineage tracing analysis of *mTmG;Nkx2.5-Cre* mice revealed both myocardial and endocardial contribution of CRE-expressing cells (Fig 5h-j), including partial recombination in the E9.5 outflow track (OFT) endocardium (Fig 5i) and in the mitral valve endocardium at E16.5 (Fig 5j). RBPJ immunostaining in E16.5 control and *Rbpj^flox^;Nkx2.5-Cre* embryos showed efficient RBPJ abrogation throughout the ventricular myocardium (99.98 ± 0.02 of cardiomyocytes recombined), while RBPJ was preserved in the majority of ventricular endocardial cells (29.06 ± 9.67% of endocardial cells recombined; Fig. 5k-l’’). In contrast, in endocardial cells overlying the valves, RBPJ depletion was significantly more efficient (76.91 ± 6.48 %; *P* <0.01 by Student’s *t*-test) (Fig. 5m-n’). Our results are in agreement with previous reports showing that NOTCH signaling abrogation by deletion of *Notch1* or *Jagged1*, using the *Nkx2.5-Cre* driver leads to VSD, DORV and BAV (18), demonstrating the requirement of endocardial

Notch signaling for valve morphogenesis, and suggesting that the lethality observed in *Rbpj^flox^;Nkx2.5-Cre* mutant mice was very likely due to *Rbpj* inactivation in valve endocardium.

These results demonstrate that myocardial inactivation of *Rbpj* in *cTnT-Cre;Rbpj^flox^* mice does not affect heart development and structure, nor does impair adult heart function, as it occurs with NOTCH signaling inactivation in the endocardium (4, 17, 18, 26, 30, 57). In contrast, *Rbpj* deletion driven by the *Nkx2.5-Cre* driver leads to VSD, DORV and BAV, phenotypes due to *Nkx2.5*-mediated CRE activity in valve endocardial cells in which RBPJ mediates NOTCH signaling, with VSD being the likely cause of perinatal lethality of these mutants.

## Conclusions

Our data indicate that: 1) Targeted inactivation of *Notch-Rbpj* in the myocardium does not affect cardiac development or function; 2) myocardial RBPJ does not mediate NOTCH signaling; 3) myocardial *Rbpj* inactivation does not affect endocardial NOTCH activity; 4) NOTCH does not play a direct role in the myocardium. Thus, the physiological role of NOTCH-RBPJ signaling in cardiac development is restricted to the endocardium, coronary endothelium and epicardium. Our data also suggest that the previously described roles for myocardial NOTCH in which the pathway was experimentally overactivated in the myocardium with various drivers (*αMhc-Cre, Nkx2.5-Cre, Mef2c-Cre*) (40, 58, 59) do not represent physiological roles for NOTCH signaling, but may offer experimental options for the manipulation of cardiac progenitors and/or lineages in the diseased heart.

## Materials and Methods

### Mouse strains and genotyping

Animal studies were approved by the CNIC Animal Experimentation Ethics Committee and by the Community of Madrid (Ref. PROEX 118/15). All animal procedures conformed to EU Directive 2010/63EU and Recommendation 2007/526/EC regarding the protection of animals used for experimental and other scientific purposes, enforces in Spanish law under Real Decreto 1201/2005. Mouse strains were *CBF:H2B-Venus* (60), *Rbpj^flox^* (42), *cTnT-Cre* (43), *Nkx2.5-Cre* (55), *Notch1^flox^* (61), mTmG (56). Details of genotyping will be provided on request.

### Tissue processing, histology and *in situ* hybridization

Embryos were fixed in 4% paraformaldehyde (PFA) at 4ºC overnight. Adult hearts were perfused with Heparin (5U/ml in PBS) and fixed during 48 hours in PFA 4%. Both embryos and adult samples were embedded in paraffin following standard protocols. Hematoxylin-eosin (H&E) staining and *in situ* hybridization (ISH) on paraffin sections were performed as described previously(62). Masson’s trichrome, Sirius Red and PAS (periodic acid-Schiff) were performed using standard procedures (CNIC Histology Facility). *mTmG;Nkx2.5-Cre* embryos were fixed in 4% PFA for an hour at room temperature, washed in PBS followed by 1 hour incubation in 30% sucrose in PBS and embedded in OCT.

### Immunohistochemistry

Paraffin sections (10 μm) were incubated overnight with primary antibodies, followed by 1h incubation with a fluorescent-dye-conjugated secondary antibody. RBPJ and N1ICD staining was performed using tyramide signal amplification (PerkinElmer NEL744B001KT). *CBF:H2B-Venus* expression was detected using anti-GFP antibody. Antibodies used in this study are: anti-RBPJ (CosmoBio 2ZRBP2, 1:50), anti-Troponin T (DSHB CT3, 1:20) anti-Cleaved Notch1 ICD (Cell Signaling Technology 2421S, 1:100), anti-GFP (Aves Labs GFP-1010, 1:400), and anti-Myosin Heavy Chain MF-20 (DHSB, 1:20). DAPI (Sigma-Aldrich D9542, 1:1000) and Isolectin B4 glycoprotein (ThermoFisher I32450, 1:100). Confocal images were obtained using Leica SP5 confocal fluorescence microscope.

### Quantification of *Rbpj* deletion

Rbpj immunostaining was analyzed using ImageJ software. Rbpj-positive nuclei were divided by the total number of nuclei (counterstained with DAPI) counted on sections both in the myocardium and endocardium of the valve and ventricles of 4 different E16.5 *Rbpj^flox^; Nkx2.5-Cre* embryos.

### Quantification of compact and trabecular myocardium thickness

H&E and images were obtained with an Olympus BX51 microscope. ImageJ software was used for the measurements, drawing a 10-pixel wide straight line along the width of compact and trabecular myocardium. Three measurements (in μm) of both compact and trabecular myocardium were taken, from both the apex and the basal region of the ventricle. Left and right ventricles were analyzed separately.

### Quantification of *Vegfa* expression

*Vegfa* expression was assessed by ISH in paraffin sections. Images of heart sections were obtained with an Olympus BX51 microscope and analyzed with ImageJ software. Using an inverted gray scale, pixel intensity was measured throughout the myocardium and a mean pixel intensity was obtained for every heart sections. 7 sections of each heart at different levels were analyzed and a mean pixel intensity was obtained for each heart.

### Ultrasound

Left ventricle (LV) function and mass were analyzed by transthoracic echocardiography in 6 months of age mice. Mice were mildly anaesthetized by inhalation of isoflurane/oxygen (1-2%/98.75%) adjusted to obtain a target heart rate of 450±50 beats/min and examined using a 30MHz transthoracic echocardiography probe. Images were obtained with Vevo 2100 (VisualSonics). From these images, cardiac output, stroke volume and LV mass were calculated. These measurements were normalized by the tibial length of each mice. Ventricular systolic function was assessed by estimating LV shortening fraction and the ejection fraction. Diastolic function was assessed by the E/A ratio. We performed a second echocardiography analysis two weeks after the first one and calculate the mean for each parameter and each mouse.

### Electrocardiograms

Electrocardiograms were recorded with and MP36 system and analyzed using the Acknowledge 4 software. 6 months old mice were anesthetized by inhalation of isoflurane/oxygen (1-2%/98.75%) adjusted to obtain a target heart rate of 450±50 beats/min.

## Statistical analysis

Statistical analysis was carried out using Prism 7 (GraphPad). All statistical test were performed using a two-sided, unpaired Student’s *t*-test. Data are represented as mean ±s.e.m. All experiments were carried out with at least three biological replicates. In the case of adult image analysis by echo and electrocardiogram analysis, the experimental groups were balanced in terms of age and sex. Animals were genotyped before the experiment and were caged together and treated in the same way. The experiments were not randomized. For adult image analysis, the investigators were blinded to allocation during experiments and outcome assessment. All quantifications are included in Supplementary Table 1.

